# DRACC1, a major postsynaptic protein, regulates the condensation of postsynaptic proteins via liquid-liquid phase separation

**DOI:** 10.1101/2023.01.23.525126

**Authors:** Takeshi Kaizuka, Taisei Hirouchi, Takeo Saneyoshi, Yasunori Hayashi, Toru Takumi

## Abstract

Numerous proteome analyses have been conducted on the postsynaptic density (PSD), a protein condensate beneath the postsynaptic membrane of excitatory synapses. Each has identified several hundred to thousands of proteins. While proteins with predictable functions have been well studied, functionally uncharacterized proteins are mostly overlooked. In this study, we perform a meta-analysis of the 35 PSD proteome datasets, including 5,869 proteins, identifying 97 uncharacterized proteins that appeared in multiple datasets. We focus on the top-ranked protein, FAM81A, renamed DRACC1. DRACC1 is expressed in forebrain neurons and enriched at the synapse. DRACC1 interacts with PSD proteins, including PSD-95, SynGAP, and NMDA receptors, and promotes liquid-liquid phase separation of those proteins. Consistently, the downregulation of DRACC1 in neurons causes a decrease in the size of PSD-95 puncta and the frequency of neuronal firing. Our results characterize DRACC1 as a novel synaptic protein facilitating the assembly of proteins within PSD. It also indicates the effectiveness of a meta-analytic approach of existing proteome datasets in identifying uncharacterized proteins.

## Introduction

Neurons communicate with each other through synapses to form a complex network. At the postsynaptic part of excitatory synapses, proteins involved in synaptic transmission and its regulation are highly enriched to form a postsynaptic density (PSD) (Dosemeci et al., 2016)(Zhu et al., 2016)(Kaizuka and Takumi, 2018). Initially identified in electron microscopic analyses of synaptic structure, the relative resistance of PSD to the detergent (Carlin et al., 1980) allowed biochemical isolation of the PSD fraction, which serves as a source for biochemical studies of the fraction (Kaizuka and Takumi, 2018). Multiple proteomic analyses of the fraction have been conducted and uncovered a number of proteins involved in synaptic transmission, including transmitter receptors, scaffolding proteins, cytoskeletal proteins, and signaling molecules (Bayés et al., 2011, 2017, 2012; Collins et al., 2006; Distler et al., 2014; Föcking et al., 2016, 2015; Han et al., 2015; Jordan et al., 2004; Li et al., 2004; Nanavati et al., 2011; Peng et al., 2004; Reim et al., 2017; Roy et al., 2018b, 2018a; Suzuki et al., 2011; Trinidad et al., 2008; Walikonis et al., 2000)(Dejanovic et al., 2018; Wilson et al., 2019). Furthermore, yeast two-hybrid screening, immunoprecipitation, affinity purification, and proximity labeling using specific PSD components pulled out direct and indirect binding partners (Dosemeci et al., 2007; Fernández et al., 2009; Li et al., 2017; Loh et al., 2016; Schwenk et al., 2012; Wilkinson and Coba, 2020).

As the technologies of proteome analysis advance, the number of proteins detected in the PSD fraction increases and reaches the order of thousands. Whereas many are *bona fide* PSD proteins from their known function and distribution, others are apparent contamination (such as proteins in glial cells or presynaptic compartments). In addition, multiple proteins without known functions or localization have been overlooked and left out of the in-depth analysis because it is difficult to determine whether they are contaminants or unreproducible.

Here, we performed meta-analyses of the multiple published datasets obtained under different experimental settings, assuming that those proteins identified in multiple datasets are authentic PSD proteins. To prove the validity of this approach, we identified and performed in-depth characterization of the top-ranked protein, Fam81, renamed as Disordered Region And Coiled-Coil Domain 1 (DRACC1). Our data confirmed that DRACC1 is indeed a functionally crucial postsynaptic molecule modulating the condensation of other PSD proteins. This approach of meta-analysis of multiple proteome datasets will help identify *bona fide* proteins in a sample that is otherwise overlooked from a single dataset.

## Results

### Meta-analysis of PSD proteome datasets

To evaluate the published PSD proteome datasets, we analyzed 35 datasets having at least 30 proteins (Table 1), of which 20 are from biochemically isolated PSD fractions followed by mass-spectrometric analysis (unbiased, Table S1, Fig. 1A) and 15 are by immunoprecipitation, pull-down, or proximal labeling of known PSD components (candidate-based, Fig. 1B). 5,869 proteins were detected at least in one dataset, where 5,800 proteins are in biochemical fractionation studies and 995 proteins in other studies (Fig. 1C). Overall, the datasets from biochemical fractionation studies showed a high overlap of identified proteins, although they are derived from various samples different from the species, brain region, and purification protocol. In contrast, immunoprecipitation, pull-down, or proximal labeling datasets showed relatively low overlap. The low overlap is observed even if the same starting protein (such as GluN2B or PSD-95) was used, suggesting the stochasticity of these approaches.

**Figure 1.**
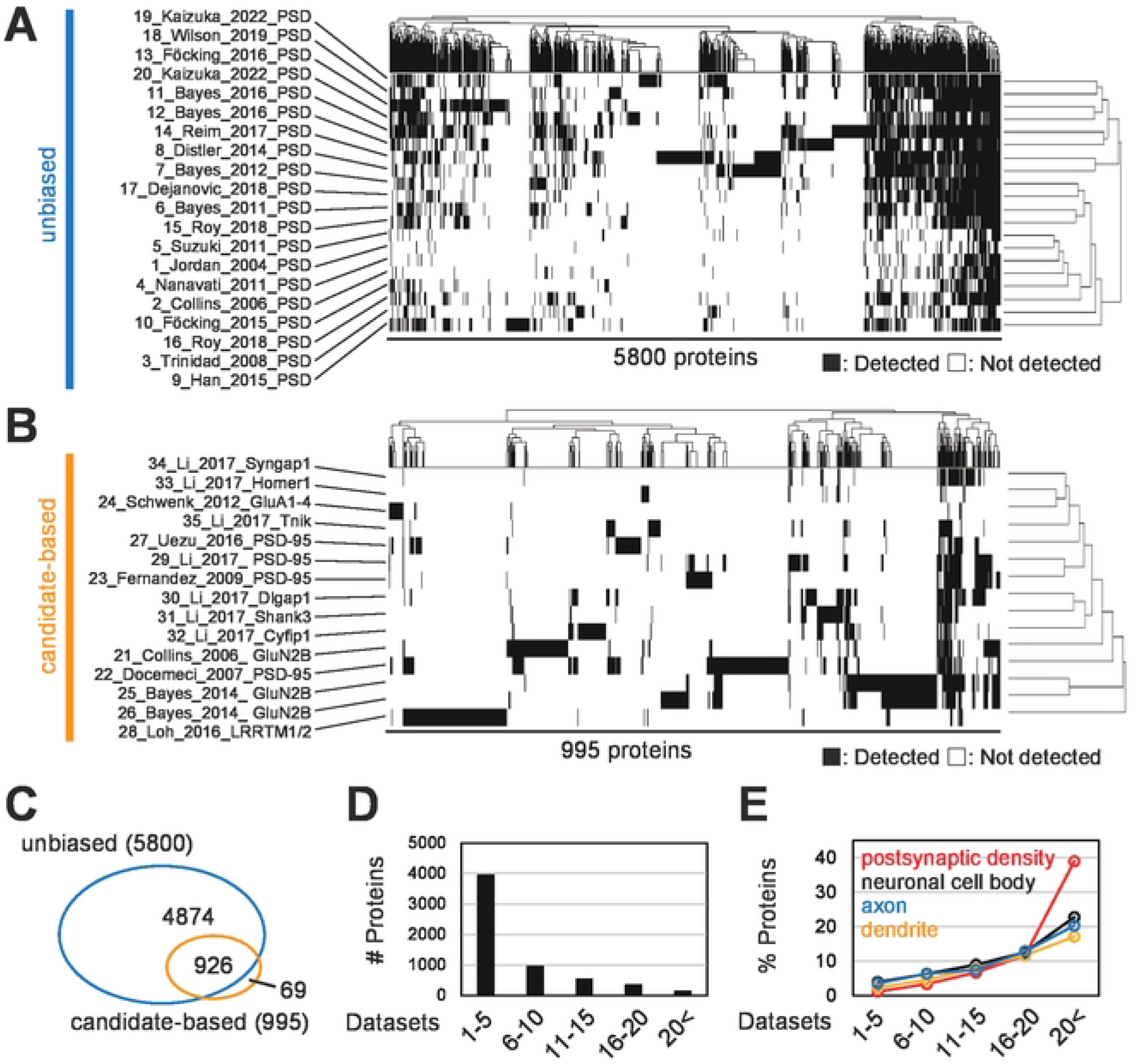
Meta-analysis of the PSD proteome datasets. (A and B) Overlap of the detected proteins in 35 PSD proteome datasets. “Protein complex” in the top panel indicates 995 proteins included in 15 datasets of the PSD protein complex. (C) Venn diagram describes the overlap of PSD fraction and PSD protein complex. (D) Histogram of the number of detected datasets. (E) The percentage of proteins that belong to indicated GO terms.

**Table 1.**
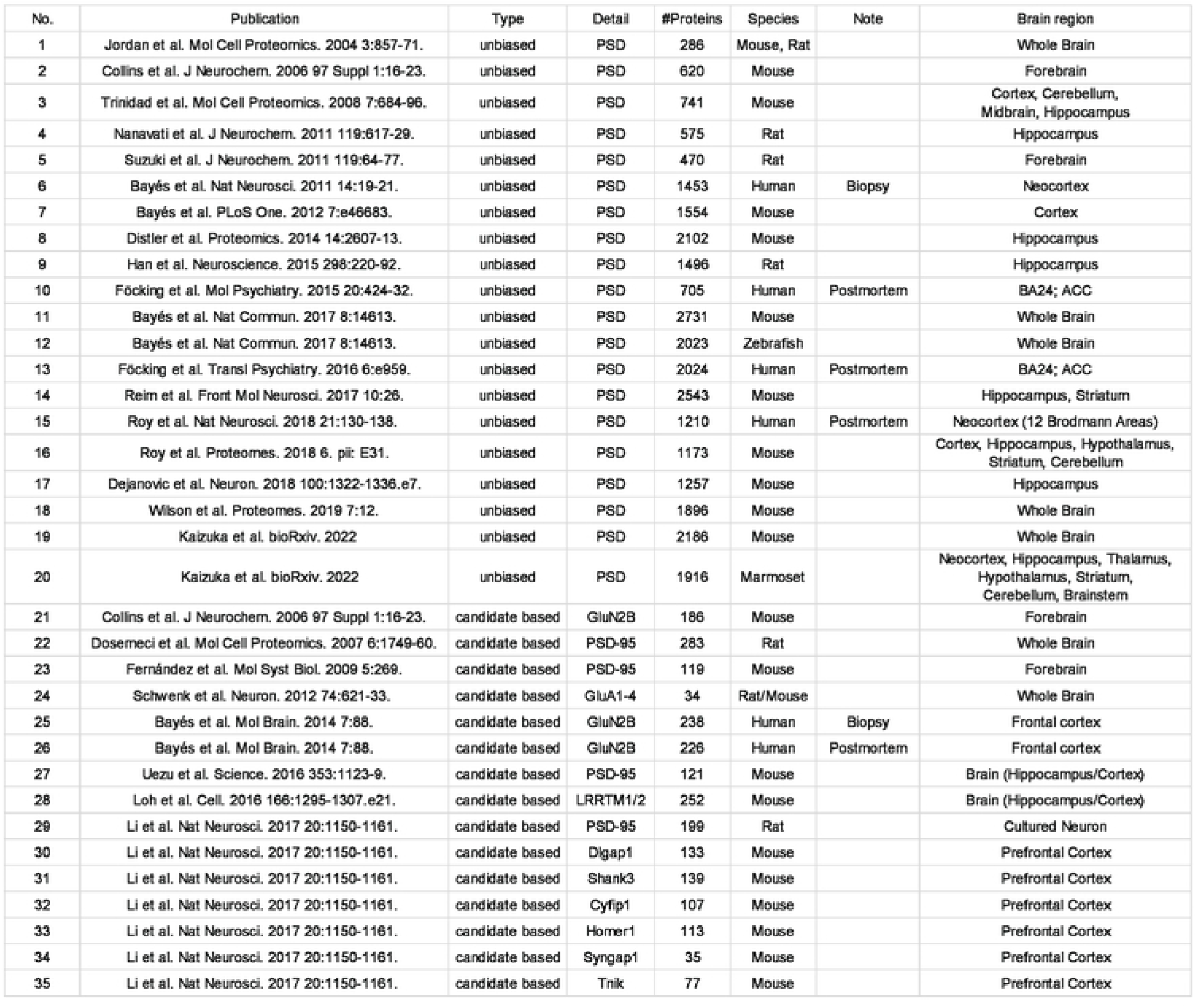
List of the 35 PSD proteome datasets referred to in this study. No.1-20 summarise the proteome data of biochemically purified PSD fraction (Fraction) and no.21-35 are that of the PSD protein complex (Complex). The original article, information on the sample and method, and the number of detected proteins of each dataset are described. Note that number of proteins can be fewer than that shown in the original article because proteins that failed ID conversion were eliminated.

Among 5,869 proteins, about 4,000 proteins were detected in only 1-5 datasets (Fig. 1D). These proteins may include contaminants from non-PSD proteins, as only 1.2% of them are known PSD proteins based on GO annotation (Fig. 1E). In contrast, proteins detected reproducibly in multiple datasets contain a higher fraction of known PSD proteins. When we analyzed 123 proteins detected in more than 20 datasets, we found that about 40% of them are PSD proteins based on GO annotation, suggesting that PSD proteins are highly enriched in this group (Fig. 1E). As expected, proteins detected in even higher (>25) number of datasets are composed of well-known core PSD proteins, including MAGUK family proteins and glutamate receptor subunits (Fig. S1). Our meta-analysis of PSD proteome datasets for the proteins recurrently detected in the multiple datasets successfully identifies known core PSD proteins.

### Identification of uncharacterized proteins from PSD proteome datasets

We asked whether this approach can detect proteins that have never been studied as PSD proteins. We first extracted protein names based on cloning ID or chromosome region (FamXX and XXXX…Rik) (Team et al., 2001; Wain et al., 2002). In addition, we also searched for proteins named after specific domains; Tmem for a transmembrane domain, Ccdc for a coiled-coil domain, Cctm for both of them, and Zfp for a zinc finger domain (Brayer and Segal, 2008; Marx et al., 2020; Priyanka and Yenugu, 2021; Schapira et al., 2017; Sohn et al., 2016). As a result, 177 proteins were identified from a total of 5,869 proteins (Table S2). Among them, 97 proteins were detected in at least 2 datasets, and 21 proteins appeared in more than 8 datasets (Fig. 2A). We focused on the top-ranked protein, FAM81A, which was detected in 21 datasets, including 15 PSD fractionations and 6 other studies (Fig. 2A and 2B). Although FAM81A is reported to localize to the PSD (Dosemeci et al., 2019), no further characterization has been reported about this protein. We, therefore, decided to focus on FAM81A to demonstrate the usefulness of the meta-analytic approach of multiple proteome databases.

**Figure 2.**
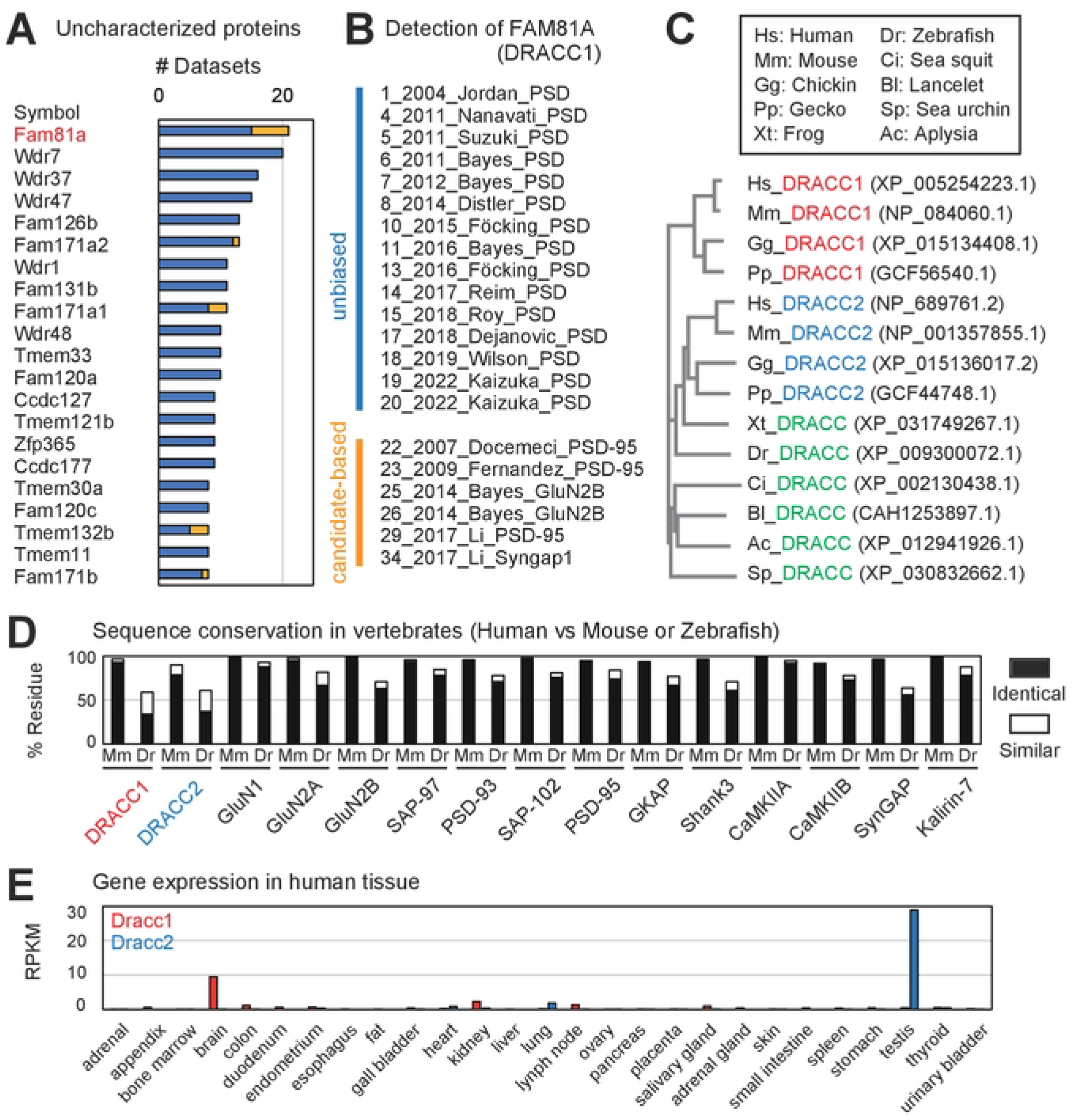
DRACC1, a PSD protein heterogeneously expressed in the higher vertebrate brain. (A) Hypothetical PSD proteins which are poorly characterized. Proteins identified in at least 8 datasets are listed. The blue and yellow bars indicate the dataset number of unbiased and candidate-based approaches. The name of proteins is described as the official symbol of the mouse gene. (B) 21 PSD proteome datasets in which FAM81A (DRACC1) was identified. (C) Homologs of DRACC1. Red and blue items indicate orthologs of DRACC1 and DRACC2, respectively. Green items indicate a common ortholog of DRACC1 and 2 in amphibians, fish, and invertebrates. The phylogenetic tree was described with Kalign (https://www.ebi.ac.uk/Tools/msa/kalign/). Hs: *Homo sapiens*, Mm: *Mus musculus*, Gg: *Gallus gallus*, Pp: *Paroedura picta*, Xl: *Xenopus laevis*, Dr: *Danio rerio*. Ci: *Ciona intestinalis*, Bl: *Branchiostoma lanceolatum*, Sp: *Strongylocentrotus purpuratus*, Ac: *Aplysia californica*. (D) Expression pattern of DRACC1 andDRACC2 in human tissues. The data was obtained from NCBI Gene. RPKM: Reads Per Kilobase of exon per Million mapped reads. (E) Sequence conservation of human DRACC1, DRACC2, and major PSD proteins in mouse (Mm) and zebrafish (Dr). The percentage of identical or similar residues was evaluated using Gene2Function.

### DRACC1 (FAM81A) is a higher vertebrate-specific PSD protein

Human FAM81A has a paralog with 35% sequence identity, FAM81B. We searched their orthologs using the protein BLAST and Gene2Function (Hu et al., 2017). Two orthologs, FAM81A and FAM81B, are found in mammals (mouse), birds (chicken), and reptiles (gecko) (Fig. 2C), whereas only one ortholog is found in amphibians (western clawed frog) and fishes (zebrafish). As for invertebrates, we found one FAM81 ortholog in tunicate (sea squirt), cephalochordate (lancelet), echinoderm (sea urchin), and mollusk (aplysia), whereas we could not find any FAM81A homologs in annelids, arthropod, and cnidaria (Fig. 2C). This suggests that FAM81 duplicated during the evolution of higher vertebrates to yield two orthologs (Zhang, 2003).

Analysis of domain architecture using the Simple Modular Architecture Research Tool (SMART) revealed that human FAM81A and FAM81B possess 2 and 4 coiled-coil domains, respectively (Fig. S2). In addition, we found that they contain regions predicted to be intrinsically disordered (Fig. S2). Considering these predictions, we named FAM81A and FAM81B as Disordered Region And Coiled-Coil Domain 1 (DRACC1) and DRACC2, respectively, and the unique orthologs in amphibians, fish, and invertebrates as DRACC.

Human DRACC1 has more than 90% sequence identity with their mouse ortholog, which is comparable to other major PSD proteins (Fig. 2D). By contrast, human DRACC1 and DRACC2 have only 34% and 37% sequence identity with zebrafish DRACC, respectively, whereas other PSD proteins have 60-80% identity with their zebrafish orthologs (Fig. 2D). These data suggest that DRACC1 evolved rapidly in the vertebrate lineage compared to other PSD proteins. Taken together, DRACC1 is a PSD protein that evolved specifically in higher vertebrates.

### DRACC1 is heterogeneously distributed in PSD within different brain regions

DRACC1 and DRACC2 are expressed explicitly in the brain and testis in humans and mice, respectively (Fig. 2E and S3A). DRACC in amphibians is expressed in a broad range of tissue, suggesting that the role of DRACC1 and 2 are differentiated during evolution (Fig. S3B). We then asked whether the abundance of DRACC1 is different across brain regions. In the mouse brain, expression of Dracc1 is limited to the forebrain and hardly detectable in the cerebellum and brainstem (Fig. S3C). In addition, the expression of Dracc1 in the cortex, hippocampus, and olfactory area is about 5-times higher than that of the striatum, pallidum, thalamus, and hypothalamus. This is in marked contrast to the global gene expression of PSD-95 (Dlg4) (Fig. S3C). Our PSD proteome data of the marmoset brain also showed region-specificity of DRACC1 expression on the PSD. PSD in the neocortex and hippocampus showed a high abundance of DRACC1 compared to other regions (Fig. S4A, S4B). PSD proteome dataset of the human neocortex shows that DRACC1 is heterogeneous across Brodmann areas (Roy et al., 2018b). The abundance of DRACC1 is relatively high in the frontal and temporal cortex compared to the parietal and occipital cortex (Fig. S4C, S4D). These data indicate that DRACC1 is heterogeneously expressed in PSD across brain regions.

### DRACC1 is enriched in dendritic spines in neurons

We then tested the subcellular localization of DRACC1. The endogenous DRACC1 was enriched in the detergent-resistant PSD fraction, similarly to the *bona fide* PSD protein, PSD-95, whereas a presynaptic protein, synaptophysin, was depleted from the fraction (Fig. 3A). In hippocampal neurons, exogenous DRACC1-GFP was colocalized with PSD-95-mCherry at dendritic spines (Fig. 3B). These results confirmed that DRACC1 is enriched in PSD, consistent with the meta-analysis of the proteome datasets as well as an earlier immunoelectron microscopic study (Dosemeci et al., 2019).

**Figure 3.**
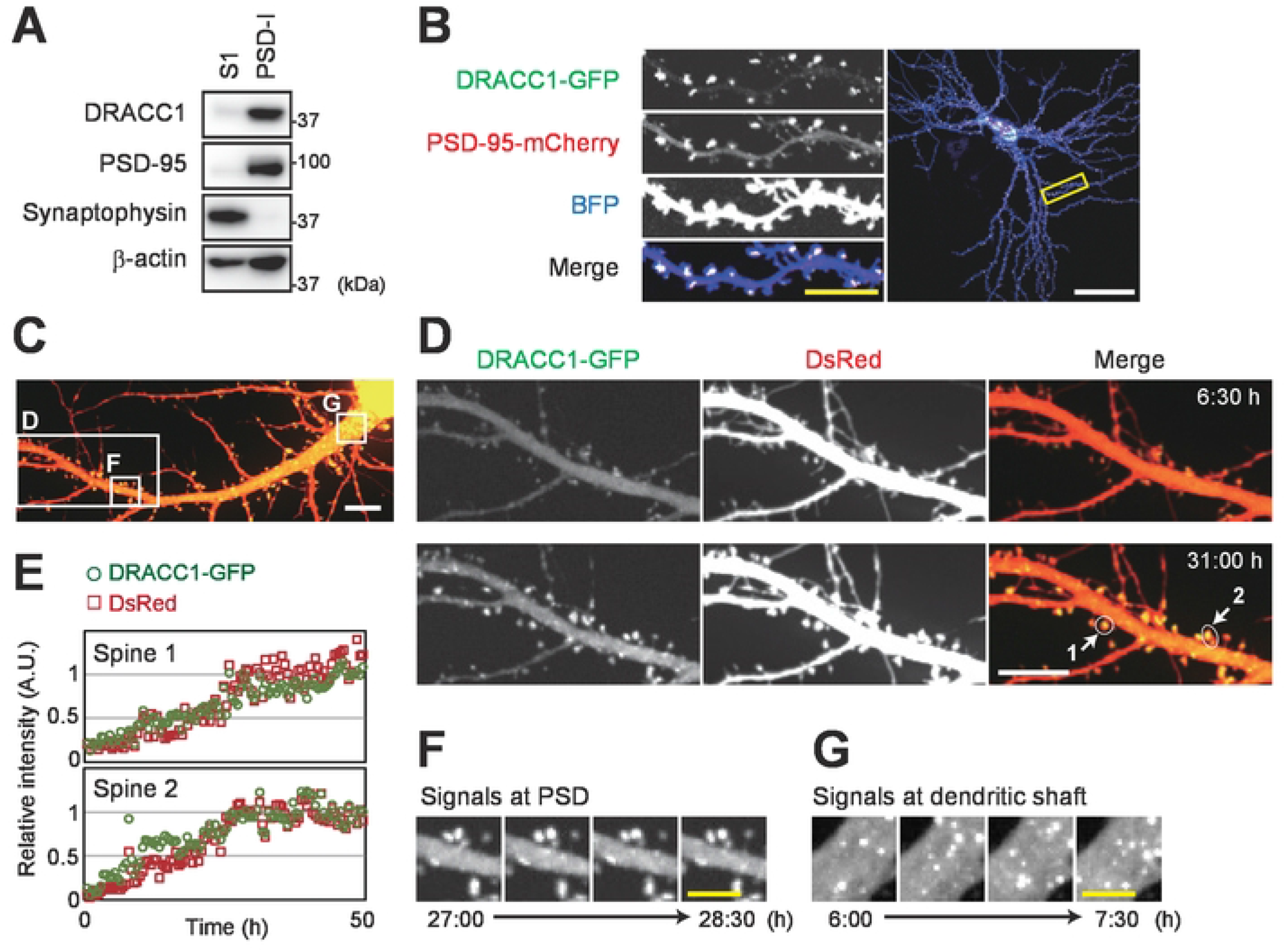
DRACC1 is a PSD protein that undergoes LLPS. (A) Enrichment of DRACC1 in PSD. S1 fraction and PSD-I fraction were prepared from brain homogenate of adult (12-week-old) mice. (B) Localization of DRACC1 at PSD in the cultured neuron. Primary mouse hippocampal neurons were transfected with DRACC1-GFP, PSD-95-mCherry, and BFP at DIV19. 2 days later, the cells were fixed and observed with a confocal microscope. Scale bar: 50 μm (white) or 10 μm (yellow). (C-G) Live-imaging of primary cultured mouse hippocampal neurons (DIV16-18) transfected with DRACC1-GFP and DsRed (30 min/frame). (C) Overview of the observed neuron. Regions magnified in panels D, F, and G are described. See also Supplementary Movie 1. (D and E) Accumulation of DRACC1 on the PSD upon spine maturation. The signal intensity of two spines shown in panel D is quantified and plotted in panel E. (F) Puncta of DRACC1-GFP on the PSD. (G) Puncta of DRACC1-GFP at dendritic shaft.

To observe the dynamics of DRACC1 in neurons, we performed time-lapse imaging of DRACC1-GFP expressed in hippocampal neurons from DIV (day in vitro) 16 to 18 (Fig. 3C, Movie S1). As the neuron matures and gains mushroom-shaped dendritic spines, DRACC1 was condensed in the structure (Fig. 3D, 3E). In addition to accumulation at dendritic spines, we found the condensates of DRACC1 in both soma and dendritic shafts (Fig. 3F, 3G). The time-lapse observation shows that the condensations are moving rapidly compared with those in PSD, which is rather stable (Fig. 3F, 3G).

### Domain structure of DRACC1 required for condensation and synaptic accumulation

We attempted to reproduce the condensate in a heterologous system and found that it can also be reproduced in HEK293T heterologous expression systems, where synaptic proteins are hardly expressed. This suggests that DRACC1 has a propensity to condensate without other synaptic molecules (Fig. 4A, Movie S2). To examine the molecular architecture of DRACC1 important for the condensation, we asked for the sequence motif required for condensation by generating a series of deletion mutants and expressing them in HEK293T cells (Fig. 4B, Fig. S5). C-terminal half (C-half; 188-364) completely abolished the puncta formation whereas N-terminal half (N-half; 1-187) still formed puncta, indicating that the N-terminal half is essential for condensation. We then made smaller deletion mutants of the N-terminal half and found that all of these mutants resulted in decreased puncta formation (Fig. 4A, 4C). In particular, Δ1-36 and Δ107-157 showed an approximately 90% decrease in the number of condensates. The size and/or maximum size of puncta of the Δ1-36 and Δ37-74 mutants were larger and more amorphous than that of full-length DRACC1, suggesting that these mutants form aggregates (Fig. 4A, 4D, 4E). Δ75-106, Δ107-157, and Δ158-187 showed a defect in the formation of enlarged structures and the size of Δ75-106- and Δ158-187-positive structures were smaller than that of full-length DRACC1, suggesting that these regions contribute to the formation of DRACC1 droplets. These results indicate that the wide-ranged sequence of DRACC1 is important for condensation.

**Figure 4.**
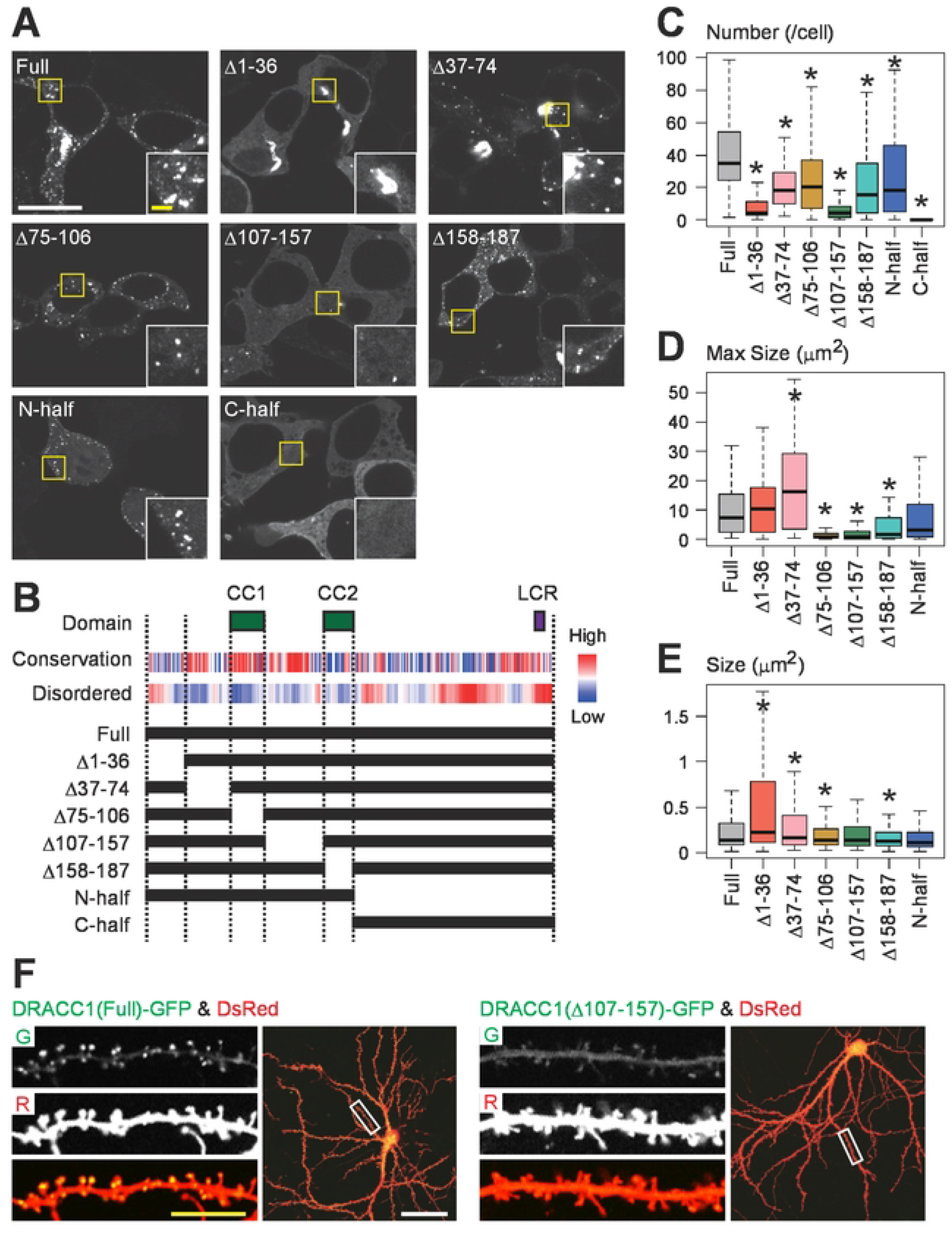
Condensation of DRACC1 is essential for its PSD localization. (A) Condensate formation of DRACC1 mutants. HEK293T cells were transfected with indicated DRACC1 mutants. 24 hrs later, the cells were fixed and observed with a confocal microscope. (B) Primary structure of mouse DRACC1. The coiled-coil domains (CC1 and CC2) and low complexity region (LCR) were identified using SMART. Evolutional conservation was analyzed using ConSurf. The probability of disorder is assessed using IUPred2A. Truncate mutants described at the bottom were used in (A). (C-E) Quantitative analysis of DRACC1 droplets described in (A). Number of DRACC1 condensates per cell (C), and size of DRACC1 condensates (D) Maximum size of DRACC1 condensates in individual cells (E) are described. The numbers of analyzed cells expressing full length, Δ1-36, Δ37-74, Δ75-106, Δ107-157, Δ158-187, N-half, or C-half DRACC1 is 98, 85, 87, 109, 95, 101, 93, 83, respectively. (F) Impaired PSD localization of the condensation deficient DRACC1 mutant. Primary cultured mouse hippocampal neurons were transfected with full-length or Δ107-157 DRACC1 mutants together with DsRed at DIV19. 2 days later, the cells were fixed and observed with a confocal microscope. Scale bars: 50 μm (white) or 10 μm (yellow). *P < 0.05

We then used the condensation-deficient mutant, Δ107-157, to test if condensation is required for synaptic accumulation in the hippocampal neuron. It showed diffused cytosolic localization and accumulation at PSD was significantly impaired compared with the full-length DRACC1 (Fig. 4F). Together, these results suggest that condensation of DRACC1 is essential for its PSD localization.

### DRACC1 forms condensates by liquid-liquid phase separation

Intriguingly, these puncta showed flexible shapes and underwent fusion and fission (Fig. 5A-5C), indicating that these punctate DRACC1 structures have liquid-like properties rather than solid aggregates. We also observed larger structures, often with complicated shapes (Fig. 5D). The shape of such structures was stable over time, suggesting that they are rigid protein aggregates (Fig. 5D, Movie S3). From these observations, we came to the idea that DRACC1 forms these droplets through the mechanism of liquid-liquid phase separation (LLPS). To test this idea, we examined the effect of 1,6-hexanediol that disrupts LLPS by interfering with hydrophobic interactions (Kroschwald et al., 2017) on DRACC1 droplets in HEK293T cells. Although large aggregate-like structures remained, most of the puncta disappeared after 10 min incubation, consistent with the idea that DRACC1 undergoes LLPS (Fig. 5E). We then tested whether the multimerization of DRACC1 is involved in LLPS. We found that GFP-DRACC1 was co-precipitated by DRACC1-FLAG expressed in HEK293T cells, suggesting that DRACC1 undergoes LLPS through multimerization (Fig. 5F).

**Figure 5.**
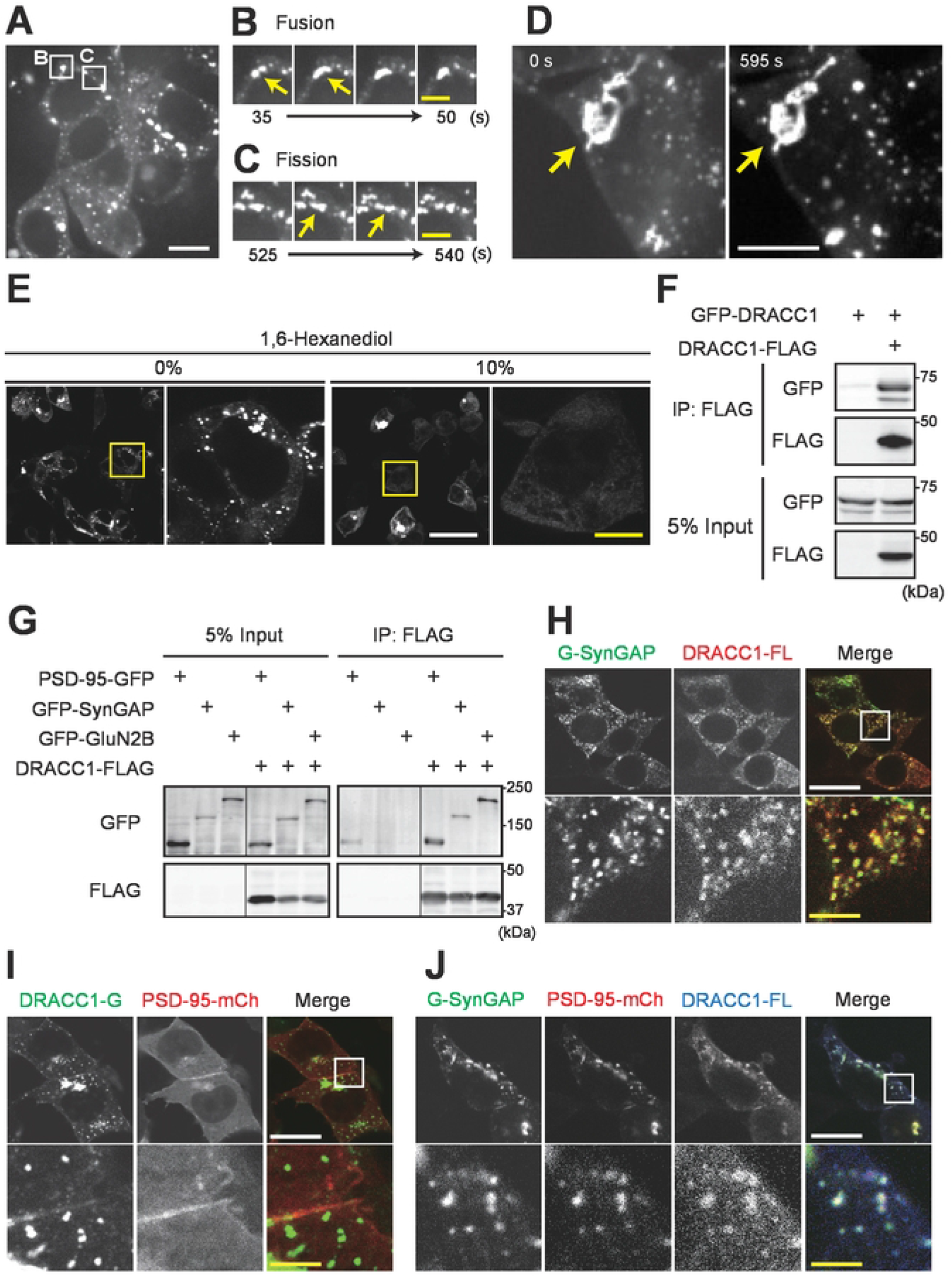
Interaction and co-localization of DRACC1 with core synaptic molecules. (A-D) Live-imaging of HEK293T cells transfected with DRACC1-GFP. (A) Overview of the observed cells. Regions magnified in panels B and C are described. See also Supplementary Movie 2. (B and C) Representative movie images of punctate structures undergoing fusion (B) and fission (C). (D) Enlarged rigid structure with a stable shape. See also Supplementary Movie 3. (E) The effect of 1,6-hexanediol on DRACC1 positive structures. HEK293T cells were transfected with DRACC1-GFP. 24 hrs later, the medium was changed to fresh medium, including the indicated concentration of 1,6-hexanediol. 10 min later, the cells were fixed. (F and G) Interaction between DRACC1 or interaction of DRACC1 and PSD-95, SynGAP, or GluN2B. HEK293T cells were transfected with indicated plasmids. 24 hrs later, cells were lysed, and immunoprecipitation was performed with an anti-FLAG antibody. Then, immunoblotting was performed with anti-GFP or anti-FLAG antibodies. (H-J) Localization of DRACC1 on SynGAP-positive droplets. HEK293T cells were transfected with indicated plasmids. 24 hrs later, cells were fixed and observed with confocal microscopy. For panels H and J, cells were subjected to immunocytochemistry using an anti-FLAG antibody before observation. Scale bars: 10 μm (white) or 5 μm (yellow) in (A-D), 50 μm (white) or 10 μm (yellow) in (E), and 20 μm (white) or 4 μm (yellow) in (H-J), respectively.

### DRACC1 interacts and forms condensate with core synaptic proteins

We then tested if DRACC1 can directly interact with the major PSD proteins. For this purpose, we first tested if DRACC1 can be co-immunoprecipitated with PSD-95, SynGAP, and GluN2B in HEK293T cells and found that these proteins indeed co-immunoprecipitate with DRACC1, indicating that they interact with DRACC1 (Fig. 5G). The interaction of DRACC1 with these proteins in nonneuronal cells, which hardly expresses synaptic molecules, suggests that DRACC1 directly binds to PSD-95, SynGAP, or GluN2B.

It has been shown that PSD-95 and SynGAP undergo LLPS when they are combined (Araki et al., 2020; Zeng et al., 2016). Consistently, we reproduced the condensation of GFP-SynGAP and PSD-95-mCherry in HEK293T cells upon co-expression (Fig. S6). Given the interaction of DRACC1 with both PSD-95 and SynGAP, we tested whether DRACC1 can undergo LLPS with SynGAP or PSD-95 in HEK293T cells. As a result, GFP-SynGAP and DRACC1-FLAG condensed together (Fig. 5H). On the other hand, PSD-95-mCherry co-expressed with DRACC1-GFP showed diffuse distribution, hardly localized with the punctate distribution of DRACC1-GFP (Fig. 5I). However, when all three were co-expressed, they formed condensate together, suggesting that PSD-95 needs SynGAP for condensation with DRACC1 (Fig. 5J). These results indicate that DRACC1 interacts with core synaptic proteins and co-localizes with SynGAP positive droplets.

To test if DRACC1 can undergo LLPS along with PSD-95, GluN2B, and SynGAP *in vitro*, we bacterially expressed and purified these proteins, combined, and observed them under a microscope. As a result, DRACC1 formed condensate in combination with SynGAP, GluN2B, and PSD-95 (3 μM each). To examine the role of DRACC1 in forming condensate, we next decreased the concentration of DRACC1 to 1 or 0 μM while maintaining the concentration of other proteins (Fig. 6). Upon reducing the concentration of DRACC1, we found a decrease in the condensate size, as visualized in the PSD-95 channel. This indicates that DRACC1 can facilitate the condensate formation of PSD proteins through the assembling and stabilizing the component proteins.

**Figure 6.**
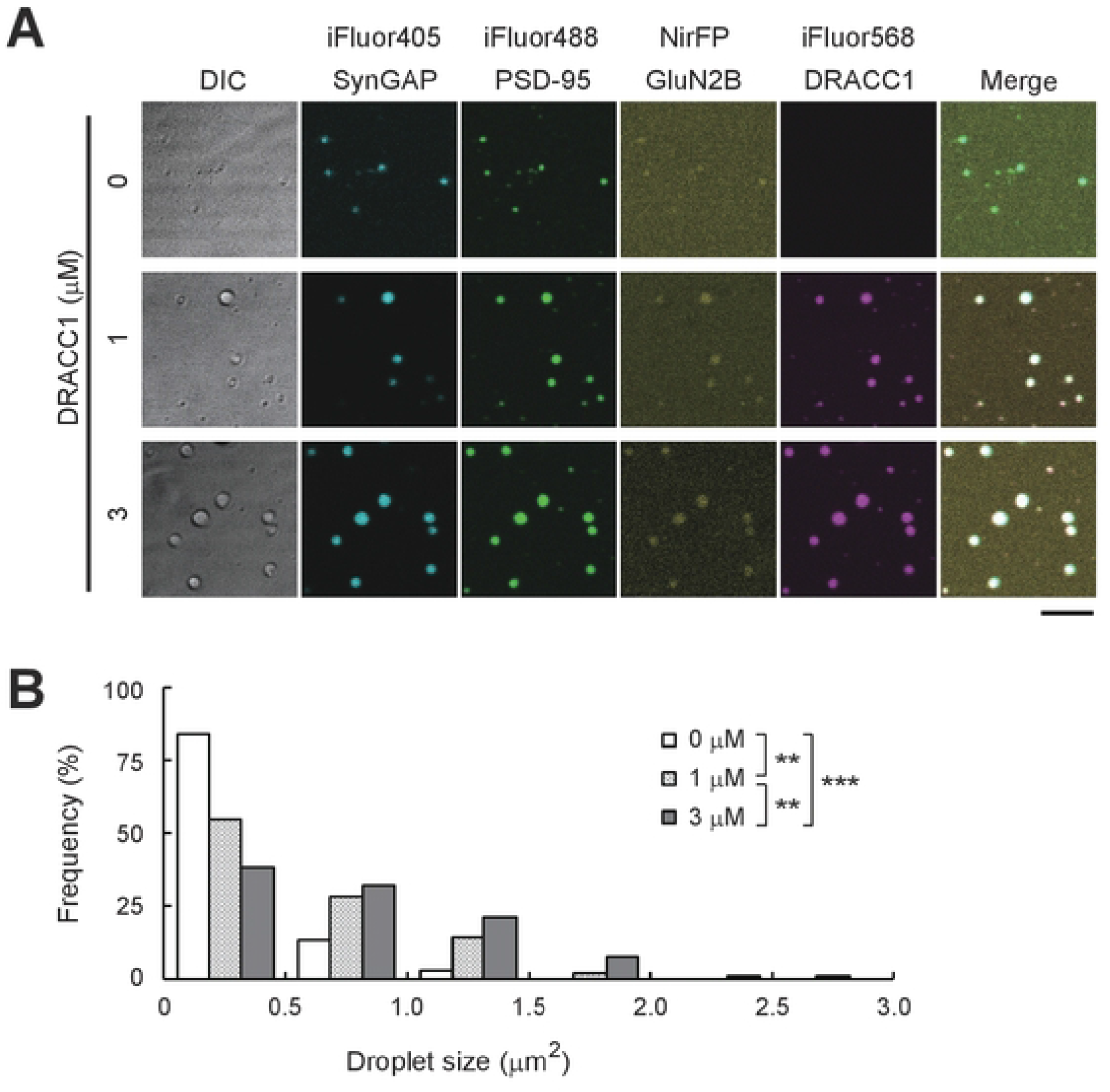
DRACC1 facilitates LLPS of postsynaptic proteins *in vitro*. (A) Confocal microscopic images of LLPS of DRACC1 with PSD-95, GluN2B carboxyl tail, and SynGAP1. iFluor 488-labeled PSD-95, NirFP-labeled GluN2B, and iFluor 405-labeled SynGAP1 (3 μM each) were mixed with increasing concentrations (0, 1, and 3 μM) of iFluor 568-labeled DRACC1. Scale bars: 5 μm. (B) The histogram of droplet size distribution. **P < 0.01, ***P < 0.001 by one-way analysis of variance (ANOVA) followed by the Tukey–Kramer test.

### DRACC1 affects PSD size and neuronal activity

To examine the role of DRACC1 on the formation of PSD in neurons, we performed a knock-down experiment of DRACC1 in the cultured hippocampal neuron. We expressed two different shRNAs against DRACC1 (shDracc1 #1 and #2) by using a lentivirus vector, both of which downregulated the mRNA level of Dracc1 to <10% (Fig. S7A). We then analyzed PSD-95 puncta on neurons, using GFP to visualize the dendrites of the infected neurons (Fig. S7B). The size of PSD-95 puncta is decreased in neurons expressing shRNA for DRACC1 (Fig. 7A, 7B), suggesting that DRACC1 stabilizes PSD-95 at the synapse. To test the physiological role of DRACC1, we next examined whether the neuronal activity is affected by DRACC1 downregulation using a multi-electrode array (MEA). We found a significant decrease in the frequency of neuronal firing in DRACC1 downregulated neurons, indicative of reduced excitatory synaptic transmission (Fig.7C, 7D). These results suggest the structural and functional importance of DRACC1 at the excitatory synapse.

**Figure 7.**
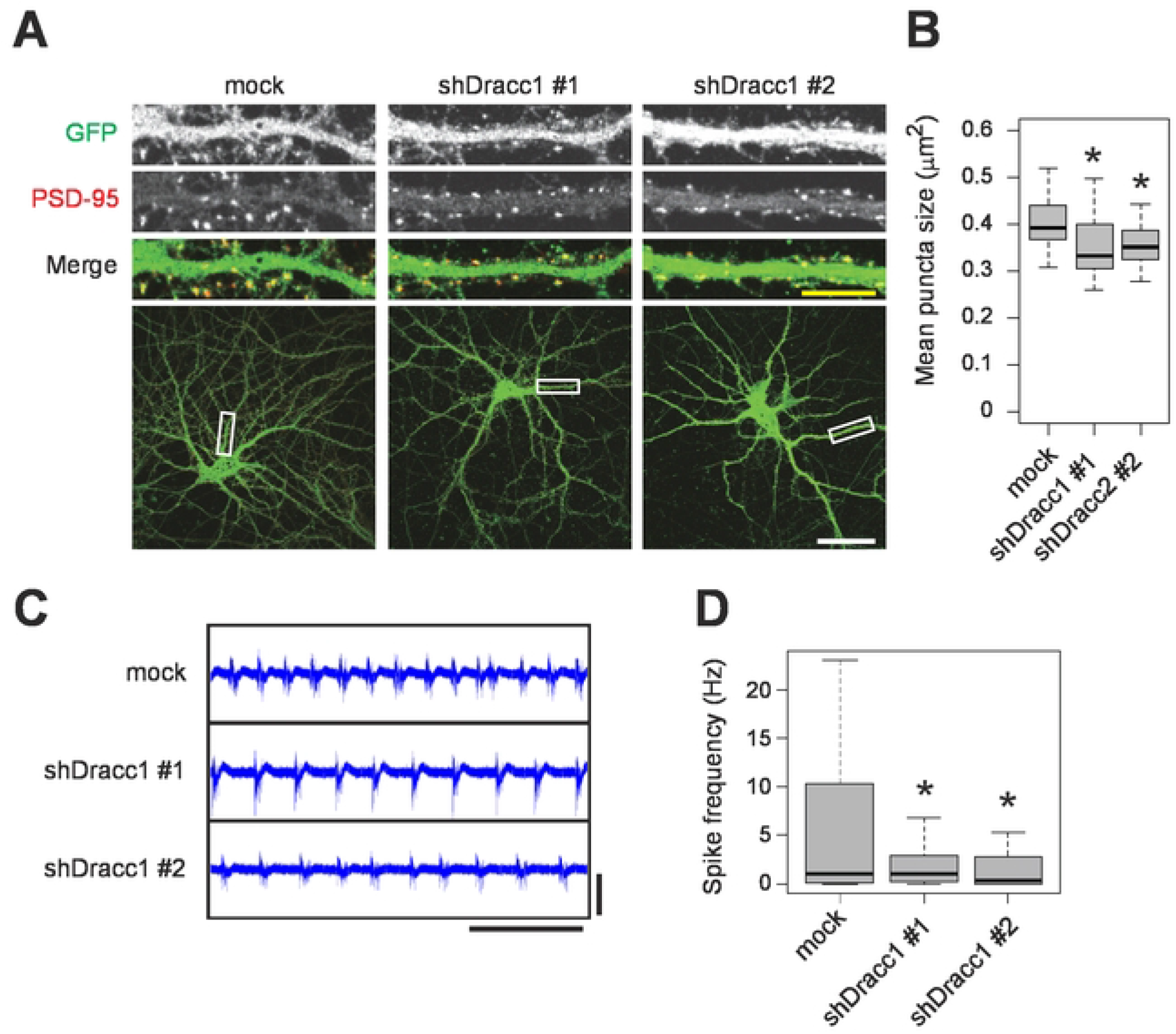
DRACC1 modulates PSD size and neuronal activity. Primary cultured mouse hippocampal neurons were infected with lentivirus-encoding DRACC1 shRNAs at DIV14. At DIV21, the neurons were subjected to immunocytochemistry (A-B) or electrophysiological recording (C-D). (A-B) Neurons were fixed and labeled with anti-PSD-95 antibody. Representative dendrite images (A) and the mean size of PSD-95 puncta (B) are described. The numbers of analyzed neurons is 51 (mock), 47 (shDracc1 #1), and 49 (shDracc1 #2), respectively. (C-D) Neurons were subjected to extracellular electrophysiological recording using MEA. Representative signals (C) and spike frequency (D) are described. Scale bars: 50 μm (A, white), 10 μm (A, yellow), 10 sec (C, x-axis), and 100 μV (C, y-axis). *P < 0.05

## Discussion

In this study, we performed a meta-analysis of the studies that identified PSD proteome and generated a versatile dataset to determine whether a given protein is in the PSD. In-depth, we characterized DRACC1, a PSD protein conserved in terrestrial vertebrates. Considering the recurrent detection of DRACC1 in the PSD proteome datasets in mammals, it is likely abundant in the PSD. Fish, amphibians, and some invertebrates have only one ortholog, DRACC. DRACC in fish is presumably not a PSD protein because DRACC was not detected in PSD proteome in zebrafish (Bayés et al., 2017), which indicates DRACC gained properties as a postsynaptic protein during evolution. We found that DRACC1 facilitates condensate formation in a dose-dependent manner. Consistently, the downregulation of DRACC1 results in decreased size of the PSD (Fig. 6, 7A, and 7B). The interaction of DRACC1 with multiple PSD proteins (Fig. 5G) and the self-interaction of DRACC1 (Fig. 5F) may be involved in the maintenance of PSD through the facilitation of LLPS between PSD molecules. The protein-protein interactions and LLPS of DRACC1 might be involved in the effect of DRACC1 on neuronal activity (Fig. 7C, 7D), as it has been proposed that synaptic activities are modulated through protein-protein interaction and LLPS in PSD (Bayer et al., 2001; Hosokawa et al., 2021; Liu et al., 2021; Saneyoshi, 2021; Saneyoshi et al., 2019). The heterogeneous expression of DRACC1 across brain regions (Fig, S3C, S4) might be involved in brain region-dependent differences in synaptic strength. Considering the limited evolutional conservation, DRACC1 may be involved in the complexity and diversity of synapses in higher vertebrates, which may be involved in the high cognitive function of these species.

DRACC1 has an intrinsically disordered domain (IDR) in its C-terminal end similarly to other proteins that undergo LLPS, such as TDP-43, FMRP, and CTTNBP2 (Shih et al., 2022; Tsang et al., 2019; Zbinden et al., 2020). Dynamics of DRACC1 in neurons observed by time-lapse imaging look similar to that of TLS/FUS (Fujii et al., 2005), which is among the first identified and best characterized RNA-binding proteins in the field of phase separation (Zbinden et al., 2020). Although we currently have no data that DRACC1 condensates contain RNA granules, including TLS/FUS, DRACC1 may play a role in signaling between synapses and dendrites. Shaft-localized condensates of DRACC1 suggest that they may regulate local translation in response to various stimuli, including synaptic transmission (Chen et al., 2020; Tsang et al., 2020) like TLS/FUS (Fujii and Takumi, 2005) or FMRP (Tsang et al., 2019) in addition to synaptic function.

A number of genetic studies point to the genes encoding excitatory synaptic proteins as causative genes for neuropsychiatric disorders such as schizophrenia and autism. We found that common variants of DRACC1 have been registered in GWAS Catalog (Buniello et al., 2019). These variants of DRACC1 (rs28890483 and rs10519005) are located at the 5’ side of the DRACC1 gene, which may affect the expression level of DRACC1. They might increase the risk of schizophrenia, bipolar disorder, and alcohol dependence. In addition, DRACC1 is reported as one of the hub genes related to susceptibility to depression (Bagot et al., 2016), suggesting that the expression level of DRACC1, which may affect neuronal activity, is involved in neuropsychiatric disorders, including major depression.

In conclusion, we provided a novel meta-analytical approach to identifying uncharacterized proteins in a given protein fraction. Using this approach, we reported an uncharacterized protein, DRACC1. The same approach will be helpful in the identification of other proteins from several existing proteomic studies on both central and peripheral tissues.

## Methods

### PSD proteome datasets and meta-analysis

Thirty-five datasets published in the following papers were referred (Bayés et al., 2011, 2014, 2012; Collins et al., 2006; Dejanovic et al., 2018; Distler et al., 2014; Dosemeci et al., 2007; Fernández et al., 2009; Föcking et al., 2015; Han et al., 2015; Jordan et al., 2004; Kaizuka et al., 2022; Li et al., 2017; Loh et al., 2016; Nanavati et al., 2011; Reim et al., 2017; Roy et al., 2018b; Schwenk et al., 2012; Suzuki et al., 2011; Trinidad et al., 2008; Uezu et al., 2016; Wilson et al., 2019). Datasets composed of less than 30 proteins were not used. The protein ID described in each report was converted to mouse Entrez Gene ID using db2db of bioDBnet (Mudunuri et al., 2009). Proteins that failed to ID conversion were eliminated from the list. Heatmaps were generated in R software using the heatmap.2 function in the gplots package.

### Mice

ICR mice were purchased from Japan SLC, Inc. All protocols for animal experiments were approved by RIKEN Brain Science Institute and Kobe University and performed by institutional guidelines and regulations. After cervical dislocation, the brain was removed from mice, briefly rinsed with ice-cold HBSS (Hanks’ Balanced Salt solution), frozen with liquid nitrogen, and stored at −80°C before use.

### Preparation of PSD fraction

Preparation of the PSD-I fraction was performed according to the previously described protocol with minor modification (Carlin et al., 1980). Briefly, three brains obtained from adult (12-week-old) ICR mice were homogenized with a glass-Teflon homogenizer in Solution A (0.32 M sucrose, 1 mM NaHCO_3_, 1 mM MgCl_2_, 0.5 mM CaCl_2_, and cOmplete EDTA-free Protease Inhibitor Cocktail). The homogenate was centrifuged at 1,400 g for 10 min at 4 °C to obtain the pellet and the supernatant. The pellet was resuspended in Solution A and centrifuged at 700 g for 10 min at 4°C. The supernatant of the first and second centrifugation was pooled as an S1 fraction and subjected to subsequent centrifugation at 13,800 g for 10 min at 4°C. The resulting pellet was resuspended with Solution B (0.32 M Sucrose and 1 mM NaHCO_3_) and centrifuged in a sucrose density gradient (0.85/1.0/1.2 M) for 2 h at 82,500 g. Synaptosomes were collected from the 1.0/1.2 M border and diluted twice with Solution B. The synaptosome was lysed by adding an equal volume of solution C (1% TX-100, 0.32 M Sucrose, 12 mM Tris-HCl pH 8.1) and rotation at 4°C for 15 min. The sample was centrifuged at 32,800 g for 20 min at 4°C to obtain PSD-I as a resulting pellet.

### PSD proteome analysis

Proteome analysis of PSD fraction obtained from mice (Dataset No.19) and common marmosets (Dataset No.20) was performed. For mouse analysis, PSD was prepared from the whole brains of 2, 3, 6, and 12-week-old ICR mice. For marmoset analysis, PSD was prepared from the neocortex, hypothalamus, thalamus, striatum, hippocampus, brainstem, and cerebellum of adult (24-month-old) marmoset. The samples were analyzed with Q Exactive HF-X mass spectrometer (Thermo Fisher Scientific) and Proteome Discoverer version 2.2 (Thermo Fisher Scientific). Proteins detected in all samples were listed and used for the meta-analysis described above.

### Plasmids

pCI-EGFP-NR2b wt (GFP-GluN2B) (Plasmid #45447), EBFP2-N1 (BFP) (Plasmid #54595), pLKO.1 - TRC cloning vector (Plasmid #10878), psPAX2 (Plasmid #12260), and pMD2.G (Plasmid #12259) were obtained from Addgene. GFP-SynGAP was gifted by Yoichi Araki and Richard Huganir (Johns Hopkins University). MG3C-SynGAP (Zeng et al., 2016) was gifted by Mingjie Zhang (Southern University of Science and Technology). pβActin empty vector, pβActin-DsRed (DsRed), and pβActin-PSD-95-GFP (PSD-95-GFP) were gifted from Shigeo Okabe (The University of Tokyo). Full-length DRACC1 cDNA (NM_029784.2) was cloned from the cDNA of the mouse brain and inserted into pEGFP-C1 (GFP-DRACC1) and pEGFP-N1 (DRACC1-GFP). To construct GST-DRACC1, DRACC1 cDNA was inserted into pGEX-6P-3. To construct DRACC1-FLAG, DRACC1 cDNA and 3xFLAG sequence were inserted into the pβActin vector. Truncated mutants of DRACC1 were constructed by inverse PCR or amplification of target sequence by PCR. To construct PSD-95-mCherry, rat PSD-95 sequence (amplified from pβActin-PSD-95-GFP) and mCherry sequence were inserted into the pβActin vector. To construct pLKO.1-GFP, the puromycin-resistant gene was removed, and the EGFP sequence was inserted into the same site. To construct shRNA plasmids, shRNA sequences were designed using Invitrogen Block-iT RNAi Designer (https://rnaidesigner.thermofisher.com/rnaiexpress/). The oligonucleotides were inserted into pLKO.1 - TRC cloning vector and pLKO.1-GFP, according to the instruction (http://www.addgene.org/protocols/plko/). Sequences of the oligonucleotides are follows; shDRACC1 #1 F: CCGGgcaactgaatcgggatattgaCTCGAGtcaatatcccgattcagttgcTTTTTG, R: AATTCAAAAAgcaactgaatcgggatattgaCTCGAGtcaatatcccgattcagttgc. shDRACC1 #2 F: CCGGgctcctggacactaaatttaaCTCGAGttaaatttagtgtccaggagcTTTTTG. R: AATTCAAAAAgctcctggacactaaatttaaCTCGAGttaaatttagtgtccaggagc.

### Cell culture

HEK293T cells and Lenti-X 293T cells (Takara, 632180) were cultured using a regular medium [Dulbecco’s Modified Eagle Medium (DMEM; Nacalai Tesque, 08458-45), 10% fetal bovine serum (FBS; Gibco, 10270), and penicillin/streptomycin (Nacalai Tesque, 26253-84)] in a 5% CO_2_ incubator. For 1,6-hexanediol treatment, a regular medium containing 10% 1,6-hexanediol (Sigma-Aldrich, 240117-50G) was used.

### Primary culture of cortical and hippocampal neurons

Hippocampi and cortices were dissected from E16.5 mouse embryos and dissociated using Neuron Dissociation Solutions (Wako, 291-78001). Neurons were counted using TC20 Automated Cell Counter (Bio-Rad) and then plated onto 24-well plates with coverslips (Matsunami, C013001), 60-mm dishes, 35-mm glass bottom dishes (Matsunami, D11130H) or MEA dishes (Multi Channel Systems, 60-6wellMEA200/30iR-Ti), which are previously coated with 0.01% poly-L-lysine in 0.1 M borate buffer solution. 4 × 10^4^ cells (for 24-well-plates), 4 × 10^4^ cells (for 60-mm dishes), 1.8 × 10^5^ cells (for 35-mm glass bottom dishes), or 1.6 × 10^5^ cells (for MEA dishes) were plated in plating media (neuron culture media plus 5% FBS). After 2-14 h, the media was changed to neuron culture media [Neurobasal (Thermo Fisher Scientific, 21103049), 1x B-27 Supplement (Thermo Fisher Scientific, 17504-044), 1x GlutaMAX-I (Thermo Fisher Scientific, 35050-061), and penicillin-streptomycin (Nacalai Tesque, 26253-84)]. At DIV4, D,L-(-)-2-amino-5-phosphovaleric acid (D,L-APV; SIGMA, A5282) was added to the media with a final concentration of 200 μM. Subsequently, half of the medium was changed once (for 24-well-late and 60-mm dishes) or twice a week (for MEA dishes).

### Live imaging of hippocampal neurons

Cells seeded on 35-mm glass bottom dishes (Matsunami, D11130H) were observed using CellVoyager CV1000 (Yokogawa Electric) equipped with a 60x objective lens. During live imaging, the culture dish was placed in a chamber to maintain incubation conditions at 37°C with 5% CO_2_. Two-color time-lapse images were acquired at 30 min or 5-sec intervals for hippocampal neurons or HEK293T cells, respectively. In the observation of hippocampal neurons, a 20 μm range of Z-stack images (21 slices, 2 μm) were acquired. As for HEK293T cells, a 2 μm range of Z-stack images (3 slices, 1 μm) were acquired. Maximum intensity projection images were shown in the figures and the movies.

### Immunoprecipitation and immunoblotting

For immunoprecipitation, HEK293T cells cultured on 10 cm dishes at ~50% confluency were transfected with 7.5 μg of DRACC1-FLAG and 7.5 μg of GFP tagged PSD-95, SynGAP, or GluN2B using PEI Max (Polysciences, 24765-1). 24 h after transfection, cells were washed with ice-cold PBS, collected with centrifugation (5,000 rpm, 2 min 4 °C), and resuspended with lysis buffer (1% Triton X-100, 50 mM Tris-HCl pH7.4, 150 mM NaCl, 1 mM EDTA, 15 mM NaF, 2.5 mM Na_3_VO_4_) with cOmplete EDTA-free Protease Inhibitor Cocktail). Lysates were kept on ice for 10 min and then centrifuged with 15,000 g for 15min at 4 °C. The supernatant was subjected to immunoprecipitation using 20 μl of ANTI-FLAG M2 Affinity Gel (Sigma-Aldrich, A2220-5ML). After 90 min incubation at 4 °C, the samples were washed five times using lysis buffer, resuspended with sample buffer (62.5 mM Tris-HCl pH 6.8, 4% sodium dodecyl sulfate (SDS), 10% glycerol, 0.008% bromophenol blue, and 25 mM dithiothreitol), and then boiled for 5 min. The samples were subjected to SDS-polyacrylamide gel electrophoresis (SDS-PAGE) and transferred onto Immobilon-FL polyvinylidene difluoride membranes (Millipore, IPFL00010). The membrane was blocked in blocking buffer (Trisbuffered saline with 0.1% Tween 20 (TBST) and 5% skim milk) at room temperature and then incubated with the indicated primary antibody overnight in blocking buffer at 4°C. The membrane was washed three times with TBST, followed by incubation with the respective secondary antibody in a blocking buffer for 1 h at room temperature. The membrane was washed five times with TBST. Immobilon Crescendo (Millipore WBLUR0500) or Chemi-Lumi One Super (Nacalai 02230-14) was used as a chemiluminescence substrate, and the signals were analyzed using ImageQuant800 (AMERSHAM). Mouse anti-FLAG antibody (F3165, SIGMA), Rabbit anti-GFP antibody (A-6455, Thermo Fisher Scientific), Rabbit anti-PSD-95 antibody (ab18258, Abcam), rabbit anti-synaptophysin antibody (#4329, Cell Signaling Technology), and mouse anti-β-actin antibody (A1978, SIGMA) were used as primary antibodies. Peroxidase goat anti-rabbit IgG (Jackson Immuno Research,111-035-003) and HRP anti-mouse IgG (Amersham, NA9310) were used as secondary antibodies.

### Immunocytochemistry and confocal fluorescence microscopy

HEK293T or hippocampal neurons grown on coverslips were washed with PBS and fixed at room temperature using 4% paraformaldehyde (PFA) in PBS for 10 min or 4% PFA and 4% sucrose in PBS for 15 min, respectively. For immunocytochemistry of endogenous PSD-95, hippocampal neurons were incubated with blocking solution [2% normal goat serum (NGS), 0.2% Triton-X-100, PBS] for 1 h at room temperature. Cells were washed with PBS twice and then incubated in antibody solution (2% NGS, 0.2% Triton-X-100, PBS) with anti-PSD-95 antibody (Millipore, MABN68) for overnight at 4 °C. Cells were washed with PBS three times and incubated with antibody solution with Alexa 568-conjugated anti-mouse IgG antibody (Molecular Probes, A-11019) for 1 h at room temperature. Cells were washed with PBS four times and embedded with Mounting Medium (Vectashield, H-1000). Cells were observed using FV3000 confocal microscopy (Olympus) equipped with a 40 × objective lens (Olympus, N2246700), a 60 × objective lens (Olympus, N1480700), or a 100 × objective lens (Olympus, N5203100). Immersion oil (Olympus, IMMOIL-F30CC) or silicone immersion oil (Olympus, SIL300CS-30CC) were used for 60 × and 100 × lenses, respectively.

### Preparation of recombinant DRACC1 protein

Preparation of recombinant DRACC1 protein was performed as follows. *Escherichia coli* strain BL21(DE3)-codonPlus-RIL (Aligent, 71136) was transformed with GST-DRACC1. Bacteria were cultured until OD600 reached ~0.8, and then isopropyl β-D-1-thiogalactopyranoside (IPTG) was added (final 0.03 mM). They were further cultured at 16°C for 18 h, centrifuged at 1,800 g for 30 min at 4°C, and then lysed with an ultrasonicator on ice (lysis buffer: 50 mM Tris, pH 8.0, 1000 mM NaCl, 10 mM imidazole, 3 mM 2-mercaptoethanol, 10% glycerol, 50 mM L-arginine, 50 mM, L-glutamic acid, 10 mM betaine, 5% trehalose and 0.2 mM PMSF). The sample was centrifuged at 100,000 g for 60 min at 4°C. GST-DRACC1 was pulled down using glutathione agarose beads (GST-accept, Nacalai-Tesque) and eluted with reduced glutathione (Nacalai Tesque, 17050-14) in the lysis buffer. While dialyzing in a buffer (25 mM Tris, pH 8.0, 100 mM NaCl, 5 mM DTT, 10% glycerol, 50 mM L-arginine, 50 mM L-glutamic acid, 10 mM betaine, 2.5% trehalose), 3C protease was added to remove GST, and then the cleaved GST tag was separated by subtractive glutathione agarose and purified in a size exclusion column buffer containing 25 mM HEPES pH 8.0, 100 mM NaCl, 0.5 mM Tris-2-carboxyethyl phosphine [TCEP] using HiLoad 26/600 Superdex 200 pg size exclusion column. Fractions (>95% purity) were pooled, concentrated, aliquoted, and flash frozen in liquid nitrogen and stored at −80°C until needed. Recombinant PSD-95, SynGAP, and GluN2B were prepared as previously described (Hosokawa et al., 2021; Zeng et al., 2016).

### Labeling and observation of LLPS of purified proteins

The DRACC1 protein was labeled by iFluor 488- or iFluor 568-succinimidyl ester (AAT Bioquest) as previously described (Hosokawa et al., 2021) The PSD proteins were diluted in a phase buffer (50 mM Tris, pH 8.0, 100 mM NaCl, 1 mM TCEP, 0.5 mM EGTA, 5 mM MgCl_2_ and 2.5 mM ATP). A protein mixture (5 μl) was injected into a homemade imaging chamber and observed by confocal microscopy (FLUOVIEW FV1200, Olympus). The number and size of the iFluor488-positive droplets were analyzed using Analyze Particles function of Fiji software.

### Lentivirus production and infection

Lenti-X 293T cells were co-transfected with pLKO.1 lentiviral plasmid, psPAX2, and pMD2.G using Lipofectamine LTX and PLUS Reagents (Thermo Fisher Scientific, 15338-100). After overnight incubation, the medium was changed to a fresh medium. 60-72 h after transfection, the supernatant was collected and filtrated through a 0.45 μm filter (Millipore, SLHV033RS). The virus was concentrated using Lenti-X Concentrator (Clontech, 631231), resuspended by one-twentieth volume of PBS, and stored at −80°C before use. For infection of cultured neurons, one-hundredth volume of virus solution was added to the neuron culture medium. 8 h after infection, the medium was changed to fresh medium. For mock infection, pLKO.1 empty vector was used as a lentiviral plasmid.

### Real-Time PCR

RNA was extracted from cortical neurons on 6 cm dishes using RNeasy Plus Mini Kit (QIAGEN, 74134) and reverse-transcribed using SuperScript II (Thermo Fisher Scientific, 18064-022). Real-time PCR was performed using the primers (F: cttagccaggctgttcttgg, R: ccagcgtctttaaggcagaa), Power SYBR Green PCR Master Mix (Thermo Fisher Scientific, 4367659), and QuantStudio3 (Thermo Fisher Scientific).

### Quantitative image analysis

1024×1024 pixel images obtained with confocal microscopy were analyzed using ImageJ. For quantification of DRACC1-GFP-positive structures in HEK293T cells, the signals were extracted by binarization using Find Maxima (noise tolerance: 120). The number and size of the signals in each cell were analyzed using Analyze Particles after selecting individual cells. Unhealthy (shrank) cells, cells with too high (saturated) or too low (invisible) signal intensity, and aggregated cells with unclear borders were avoided from the analysis. Images were first shuffled for blinded analysis to quantify PSD-95-positive structures in neurons expressing GFP. Then, images with single neurons were selected. PSD-95 positive structures were extracted by binarization using Find Maxima (noise tolerance: 120). Structures with more than 50 pixels were eliminated to avoid the detention of non-synaptic structures. To assess dendrite length, signals of GFP were traced using NeuronJ (Pool et al., 2008). The PSD-95 puncta along the dendrites (within 25-pixel diameter) were extracted using SynapCountJ (Mata et al., 2016). The mean puncta size and the number of puncta per dendrite length were calculated for individual neurons. The result was visualized as a boxplot using R software. Student t-test was used for statistical analysis.

### Multi-electrode array (MEA)

The medium was changed with a fresh neuron culture medium without D,L-APV. 30 min after the medium change, neuronal activity was analyzed using MEA2100 (Multi Channel Systems) and MC_Rack Version 4.6.2. Input voltage range and sampling frequency were set as ±19.5 mV and 20000 Hz, respectively. Neuronal activity was recorded for 2 min. Voltage over 5 standard deviations was used as a threshold to detect spikes.

### Databases and bioinformatics tools

Metascape (https://metascape.org/gp/index.html#/main/step1) (Zhou et al., 2019) was used to analyze the protein-protein interaction network. NCBI Gene (https://www.ncbi.nlm.nih.gov/gene/) was referred to check gene expression in humans and mice. Xenbase (http://www.xenbase.org/entry/) (Fortriede et al., 2020) was referred to check gene expression patterns in the frog. Protein BLAST (https://blast.ncbi.nlm.nih.gov/Blast.cgi) was used to identify orthologs of proteins. Gene2Function (https://www.gene2function.org/) (Hu et al., 2017) was also referred to find orthologs in invertebrates and unicellular organisms. Allen Brain Atlas (http://mouse.brain-map.org/) was referred to check gene expression patterns in mouse brains. SMART (http://smart.embl-heidelberg.de/) (Letunic and Bork, 2018) was used to analyze domain architecture. D2P2 (http://d2p2.pro/) (Oates et al., 2013) was used to evaluate disordered regions. ConSurf (Ashkenazy et al., 2016) was used to assess evolutional conservation. MARRVEL (http://marrvel.org/) (Wang et al., 2017) was referred to search for rare variants. GWAS Catalog (https://www.ebi.ac.uk/gwas/home) (Buniello et al., 2019) was referred to search common variants.

## Author contributions

T.K. designed the research and performed most experiments and data analysis. T.H. and T.S. performed *in vitro* liquid-liquid phase separation experiment. T.K. wrote, and Y.H. and T.T. edited the manuscript. Y.H. and T.T. supervised the work. All authors commented on the manuscript and approved the final version.

## Acknowledgments

We thank Y. Araki, R. Huganir, M. Zhang and S. Okabe for providing the plasmids and A. Okuda, K. Morishima, M. Sugiyama, T. Imasaki, and R. Nitta for helpful discussion. We thank T. Otani, C. Noguchi, S. Yoshida, and S. Fujima for their technical assistance and all technical staff in the Takumi lab for preparing experimental reagents. We thank Jessica Griffiths for proofreading the manuscript. This work was supported in part by KAKENHI (JP16H06316, JP16H06463, JP18K14830, JP18H05434, JP20K21462, JP21H00202, JP21H04813, JP21K19351, JP21H02595, JP21H05692) from the Japan Society for the Promotion of Science (JSPS) and the Ministry of Education, Culture, Sports, Science, and Technology (MEXT), Japan Agency for Medical Research and Development (AMED) under Grant Number JP21wm0425011, Japan Science and Technology Agency (JST) under Grant Number JPMJMS2299 and JPMJCR20E4, Intramural Research Grant (30-9) for Neurological and Psychiatric Disorders of NCNP, the Takeda Science Foundation, Smoking Research Foundation, Tokyo Biochemical Research Foundation, Taiju Life Social Welfare Foundation, The Naito Foundation, and The Tokumori Yasumoto Memorial Trust for Researches on Tuberous Sclerosis Complex and Related Rare Neurological Diseases, and Human Frontier Science Program (RGP0020/2019). T.K. was supported by Grant-in-Aid for JSPS Fellows (JP16J04376).

---

**Figure S1. Proteins detected in multiple datasets include core PSD proteins**

List of the top 27 proteins that show the highest number of datasets. Blue and yellow bars indicate the dataset number of unbiased and candidate-based approaches. Proteins detected in at least 26 datasets are listed.

**Figure S2. Domain architecture of FAM81A/DRACC1 homologs**

Coiled-coil regions detected by SMART and disordered regions predicted by IUPred2A.

**Figure S3. Gene expression pattern of DRACC1 and DRACC2 in mouse and frog**

(A and B) Gene expression pattern of Dracc1 and Dracc2 in mouse (A) and frog (B) tissue. Mouse data and frog data were obtained from NCBI Gene and Xenbase, respectively. (C) Gene expression pattern of Dracc1 and Dlg4 (PSD-95) in mouse brain. The data was obtained from Allen Brain Atlas (Experiment 73732150 and 72109445). RPKM: Reads Per Kilobase of exon per Million mapped reads, TPM: Transcripts Per Kilobase Million.

**Figure S4. DRACC1 is differentially expressed across brain regions**

(A and B) The abundance of DRACC1 on PSD across different brain regions in the common marmoset. (A) Graphical illustration of brain regions analyzed in the dataset. (B) The plot of relative abundance of DRACC1. Data from three technical replicates are shown. Green lines indicate the average. (C and D) The abundance of DRACC1 on PSD across Brodmann areas (BAs) in the human brain (Roy et al., 2018b). (C) Graphical illustration of BAs analyzed in the dataset. The red to blue color gradient indicates the mean relative abundance of DRACC1. (D) Plot of the relative abundance of DRACC1. Data from four individuals (Brain A, B, C, and D) are shown. Green lines indicate the average relative abundance.

**Figure S5. Protein sequences of DRACC1 orthologs in mammals, birds, and reptiles**

The sequence of DRACC1 homologs of indicated species are aligned using Kalign and visualized using MView. The numbers on the sequences indicate the amino acid number of mouse DRACC1. Residues identical to human DRACC1 are highlighted. The colors of the characters represent the classification of amino acids; hydrophobic (light green), large hydrophobic (dark green), positive (red), small alcohol (light blue), and polar (purple). Mammals: human (*Homo sapiens*), rat (*Rattus norvegicus*), and mouse (*Mus musculus*). Birds: chicken (*Gallus gallus*) and zebra finch (*Taeniopygia guttata*). Reptiles: soft-shelled turtle (*Pelodiscus sinensis*) and gecko (*Paroedura picta*). The coarse-coil domains (CC1 and CC2) and low complexity region (LCR) of mouse DRACC1 are labeled with a black bar.

**Figure S6. Condensates of PSD-95 and SynGAP**

HEK293T cells were transfected with indicated plasmids. 24 hrs later, cells were fixed and observed with confocal microscopy. Scale bars: 20 μm (white) or 4 μm (yellow).

**Figure S7. Knockdown of DRACC1 with lentivirus shRNA**

(A) Primary cultured mouse cortical neurons were infected with lentivirus encoding DRACC1 shRNAs at DIV4. At DIV16, neurons were harvested to extract mRNA. After the preparation of cDNA, real-time PCR was performed. The relative expression level is described. (B) Primary cultured mouse hippocampal neurons were infected with lentivirus of pLKO.1-GFP mock plasmid at DIV14. At DIV21, the neurons were fixed and subjected to immunocytochemistry using an anti-PSD-95 antibody. Scale bars: 100 μm.

**Table S1. List of 5,869 proteins included in PSD proteome datasets**

Proteins were described as mouse Entrez Gene ID. 0 and 1 indicate undetected and detected, respectively.

**Table S2. List of 177 proteins in PSD proteome datasets that have not been fully characterized**

**Supplementary Movie 1. DRACC1 droplets in hippocampal neuron**

Mouse primary hippocampal neurons were transfected with DRACC1-GFP and DsRed. Images were obtained every 30 min.

**Supplementary Movie 2. DRACC1 droplets in HEK293T cells**

HEK293T cells were transfected with DRACC1-GFP. Images were obtained every 5 seconds.

**Supplementary Movie 3. Aggregation-like structure of DRACC1 in HEK293T cells**

HEK293T cells were transfected with DRACC1-GFP. Images were obtained every 5 seconds.

